# Ponds, puddles, floodplains and dams in the Upper Xingu Basin: could we be witnessing the ‘lentification’ of deforested Amazonia?

**DOI:** 10.1101/524652

**Authors:** Luis Schiesari, Paulo R. Ilha, Daniel Din Betin Negri, Paulo Inácio Prado, Britta Grillitsch

**Affiliations:** Gestão Ambiental, Escola de Artes, Ciências e Humanidades, Universidade de São Paulo, São Paulo, SP, 03828-000, Brazil; Departamento de Ecologia, Instituto de Biociências, Universidade de São Paulo, São Paulo, SP, 05508-090, Brazil; Instituto de Pesquisa Ambiental da Amazônia, Canarana, MT, 78640-000, Brazil; LAGE at Departamento de Ecologia, Instituto de Biociências, Universidade de São Paulo, São Paulo, SP, 05508-090, Brazil; Department of Biomedical Sciences, University of Veterinary Medicine of Vienna, Vienna, 1210, Austria

**Keywords:** land use change, deforestation, soybean, pasture, dams, freshwater, fish, amphibians

## Abstract

Hydrological change is a conspicuous signal of land use intensification in human-dominated landscapes. We hypothesized that land conversion and land use change increase the availability of lentic habitats and associated biodiversity in Southern Amazonian landscapes through at least four drivers. River damming promotes the formation of reservoirs, which are novel permanent lentic water bodies. A rise in the water table driven by local deforestation promotes the expansion of shallow riparian floodplains. Soil compaction and the deliberate construction of cattle and drainage ponds promote the increase in temporary water bodies in interfluvia. We tested these hypotheses using data on habitat characterization and biological surveys of amphibians and fish in forests, pastures and soybean fields in the headwaters of the Xingu River in Mato Grosso, Brazil. Lentic habitat availability sharply increased in deforested land, with consequences to freshwater biodiversity. Reservoir formation influenced both fish and amphibian assemblage structure. Fish species ranged from strongly favored to strongly disfavored by reservoir conditions. Amphibian richness and abundance increased in pasture and soybean streams relative to forests, with a strong positive effect of density of reservoirs in the landscape. Expansion of stream floodplains increased the abundance of *Melanorivulus megaroni*, a fish species indicator of shallow lentic habitats. Rainwater accumulation in temporary ponds and puddles, entirely absent from well-drained forested interfluvia, allowed the invasion of converted interfluvia by twelve species of open-area amphibians. A literature review indicates that these four drivers of hydrological change are geographically widespread suggesting that we may be witnessing a major yet previously unaccounted form of habitat change in deforested Amazonia.

## Introduction

Hydrological change is a conspicuous signal of land use intensification in human-dominated landscapes [1]. The importance of water as a resource for human, livestock and crops, as a source of energy, and as pathway for transportation makes water management a quintessential element of human activities. In addition to the deliberate management of waterbodies, interventions in the terrestrial environment indirectly influence hydrology by altering the partitioning of rainfall among the processes of evapotranspiration, infiltration and runoff [2].

Frontier landscapes provide a relevant environmental scenario for testing the occurrence and consequences of anthropogenic hydrological change both because ongoing land use change permits side-by-side comparison of converted and native habitats, and because frontiers are regions where biodiversity change is expected to be highest [3]. The Amazon Basin is the largest area of the world currently undergoing frontier settlement and, within the Amazon Basin, the Brazilian state of Mato Grosso accounts for more than 40% of deforested land [4]. Building on published hydrological studies [5; 6; 7; 8], we test the hypothesis that deforestation is promoting the ‘lentification’ of Southern Amazonian landscapes, with consequent shifts in their freshwater fauna. The term ‘lentification’ was coined to the natural lentic characteristics of small, intermittent floodplain channels during low waters [9] and later used to the transformation of the habitat character of rivers from lotic (i.e. flowing water) to lentic (i.e. standing water) through processes such as flow regulation, unsustainable water extraction and climate change [10; 11]. The term does not appear to have been embraced by the scientific community though (only three papers with ‘lentification’ were retrieved in a Web of Science search; March 13, 2018) and lacks an operational definition. We here define ‘lentification’ as an increase in the availability (i.e. density and/or area cover) of lentic freshwater habitats in the landscape, directly or indirectly caused by human activities. This definition is inclusive of any driver, and highlights that lentification is a property of the landscape and not only of the drainage network as it includes the creation and change of freshwater habitat in interfluves as well. Because few processes lead to the formation of novel lotic habitats, an increase in the availability of lentic habitat almost invariably implies an increase in the proportion of lentic habitats in the landscape.

We hypothesize that at least four drivers of lentification could be operating in the Upper Xingu Basin, one of the largest tributaries of the Amazon river: stream and river damming, increased water yields, soil compaction, and the deliberate construction of cattle and drainage ponds. Among these, river damming is the most readily recognized driver of lentification as the Amazon Basin is the new ‘hydroelectric frontier’ in Brazil with 154 hydroelectric dams in operation, 21 under construction and 277 planned [12]. Less attention is given to the myriad of small dams found today in deforested land. Yet, in the Upper Xingu Basin alone there are ~10,000 small dams, most of which were built to provide water to cattle and generate electricity for local consumption [6]. Lentification is a clear product of river damming as reservoirs have attributes typical of lentic systems including slow currents, high water residence times, greater depth and surface area, open canopy, high surface temperatures, thermal stratification, and net accumulation of sediment and nutrients [13].

Deforestation could also promote lentification through increased water yields. Local deforestation has little effect on precipitation but strongly reduces per-unit-area evapotranspiration due to the lower height, shallower rooting depth, and lower leaf area index of grasses and crops relative to trees. Thus, proportionally more rainfall becomes river discharge [2]. In the Upper Xingu Basin, mean daily water yields increased four times in soybean watersheds relative to forest watersheds via an increased contribution of groundwater-driven baseflow to stream discharge [7]. Lentification could result if deforestation-driven increased water yields promote the expansion of semi- lentic riparian floodplains. Note, however, that this effect is strongly scale-dependent. When deforestation reaches regional scales, the reduction in evapotranspiration feeds back into reduced rainfall and therefore, reduced water yields [e.g.,14].

In interfluves away from streams and rivers, deforestation could also promote lentification via soil compaction. In the Upper Xingu Basin soil infiltrability was markedly reduced in the conversion of forests to pastures both in terms of infiltrability (from 1258 mm/h to 100 mm/h) and saturated hydraulic conductivity, probably due to cattle trampling [5]. Shallow disking in the conversion of pastures to soybean fields increased infiltrability (469 mm/h) but further reduced subsoil saturated hydraulic conductivity [5]. Lentification could result if deforestation-driven soil compaction increases the frequency in which precipitation exceeds infiltration, thereby forming ponds and puddles. An additional driver of lentification in interfluvia is the deliberate construction of ‘cacimbas’, artificial ponds providing water to cattle and draining stormwater runoff from dirt roads.

Taken together, it is apparent that there are several drivers in place capable of creating novel lentic water bodies in converted Amazonian landscapes, and that such novel lentic water bodies span the full spectrum of the hydroperiod gradient – from ephemeral to semi-permanent to permanent waterbodies – in such a way that could consistently favor quite distinct elements of lentic freshwater fauna [15].

In this study we test, for the first time, the hypotheses (i) that land conversion in the Upper Xingu Basin is associated with an increase in the availability of lentic freshwater habitats both along drainage networks (by creating reservoirs and expanding stream floodplains) and in interfluves (by creating ponds and puddles), and (ii) that this landscape-level lentification increases the abundance of freshwater fauna typically associated with lentic habitats. We tested these hypotheses using data on habitat characterization and biological surveys of fish and amphibians in forests, pastures and soybean fields in the headwaters of the Xingu River in Mato Grosso, Brazil. Fish and amphibians are excellent indicators of hydrological change [16] with high complementarity in habitat use [15]: fish are usually restricted to permanent waterbodies whereas most amphibians reproduce in temporary and semi-permanent lentic waterbodies. Importantly, our study was conducted at the same site where the hydrological mechanisms we envision as promoting lentification were demonstrated. Hayhoe et al.[7] demonstrated that deforestation increases water yields but not whether this phenomenon increases the area of shallow riparian floodplains. Scheffler et al.[5] demonstrated that deforestation decreases soil infiltrability but not whether this phenomenon promotes the formation of ponds and puddles. Finally, Macedo et al.[6] quantified the number of impoundments in the Upper Xingu Basin as a whole; we here confirm their findings at our specific study location before linking habitat with biodiversity change.

## Methods

### Study area

This study was conducted in the headwaters of the Xingu River in Canarana and Querência, Mato Grosso, Brazil (Fig 1). Climate is Köppen’s ‘Aw’, a tropical savanna climate with distinct wet (October-April) and dry (May-September) seasons. Mean annual rainfall is 1898 mm (1987-2007; Grupo AMaggi, pers.comm.). Landscape is dominated by wide interfluves where plateaus 27-62 m high gently slope towards stream channels. Water table is ~20-40 m deep. Soils are ustic oxisols (‘latossolos vermelho-amarelos distróficos’ according to the Brazilian classification) in plateaus grading into aquic inceptisols (‘gleysolos’) in riparian zones [7]. Original vegetation cover is a closed-canopy, evergreen seasonal forest that is transitional between the ombrophilous rainforests in the north and the cerrados in the south of the Xingu Basin [17].

**Fig 1.**
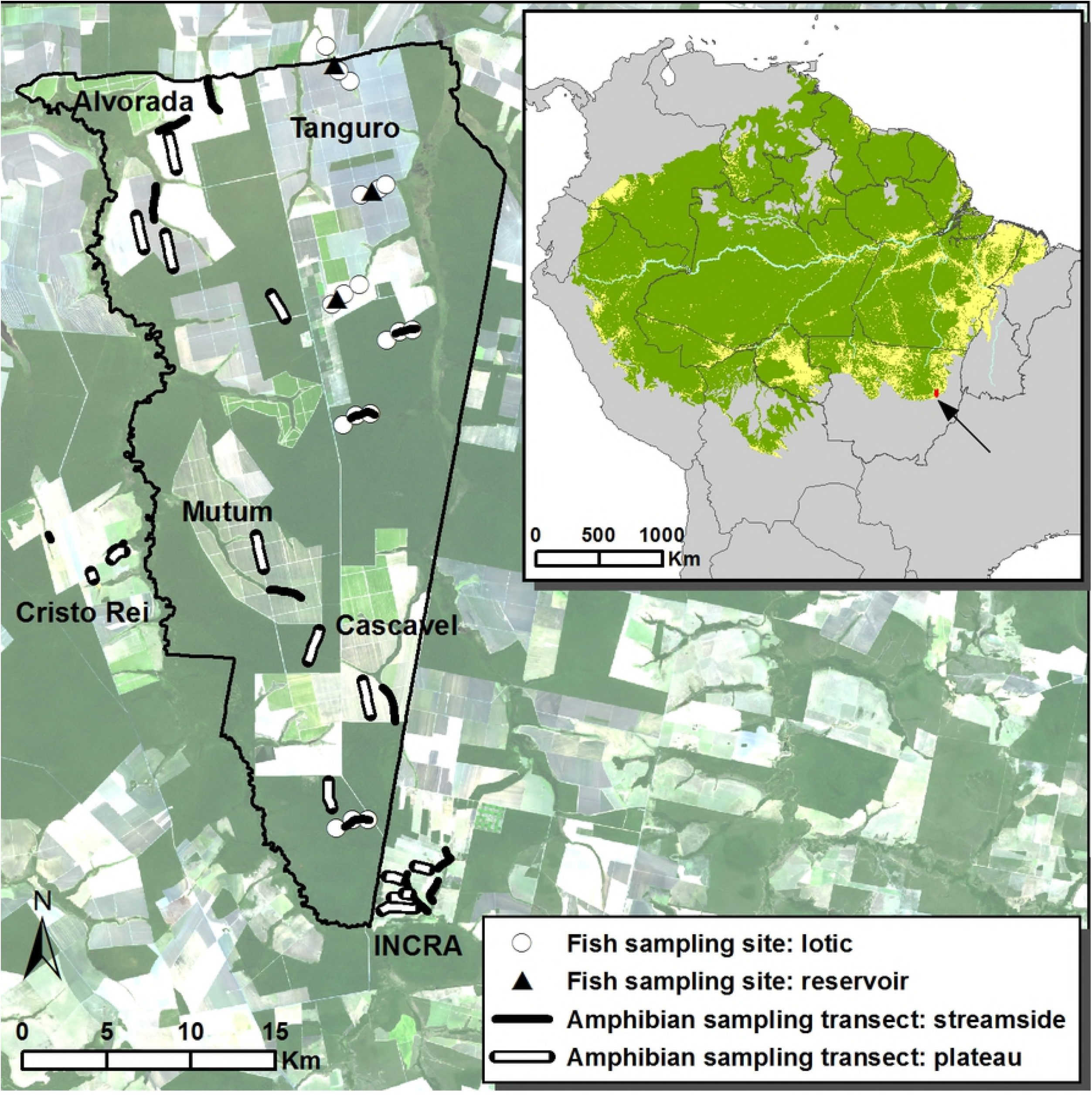
Fish sampling sites and amphibian sampling transects in Tanguro Ranch and surrounding farms (Landsat 8; July 2, 2013). Dark green represents primary closed canopy forests, light green represents deforested areas (soybean fields in Tanguro Ranch, pastures in Cristo Rei Ranch and in the INCRA settlement). From North to South, streams sampled for fish are Tanguro C, Tanguro B, Tanguro A, APP2, APP2A, and APPM; transects in forest plateaus are Seringal, AU and A1. Not represented is Lírio Branco Ranch, the third replicate pasture, located 27 km to the South. The inset shows the location of the study site in Southeastern Amazon. In the Amazon Basin, green represents closed-canopy forests, yellow represents deforested areas [according to 18] and uncolored areas represent native savannas. Savannas outside of the Amazon Basin are not represented but dominate the original vegetation cover a few tens of kilometers South and East of the study site. Note the position of the study site in the agricultural frontier known as ‘the Amazonian Arc of Deforestation’.

Fieldwork was conducted in and around Tanguro Ranch (Fig 1), a 82,000 ha farm covered with primary closed-canopy forests (44.000 ha) and soybean plantations (38,000 ha). Neighboring Cristo Rei Ranch, Lírio Branco Ranch and the INCRA Settlement were predominantly covered by pastures. At the time of sampling streams in deforested watersheds were by law bordered by a 25-m wide riparian forest buffer, which structural integrity was variable across the region.

### Testing for a land use-driven lentification of the landscape

#### Density of impoundments in streams

To test the hypothesis that land conversion is associated with an increase in the density of man-made impoundments, we defined one stream transect in each of three forests (along APPM, APP2 and APP2A streams in Tanguro Ranch), three pastures (INCRA Settlement, Cristo Rei Ranch, Lírio Branco Ranch) and three soybean fields (Alvorada, Mutum and Cascavel fields in Tanguro Ranch) (Fig 1). One pasture (INCRA Settlement) and one soybean field (Alvorada) received two additional transects each for a total of 13 transects. These ‘focal’ pasture and soybean fields were selected for detailed physical, chemical and biological sampling surveys (to be published elsewhere). We could not select a ‘focal’ forest with replicated transects due to the scarcity of forest trails. Each transect was 2000 m long, and positioned at least 400 m away from other transects or from other land uses. Using these criteria, transect position was randomly defined among all available possibilities. In February 2012 we recorded the number of impoundments per stream transect.

#### Density of ponds and puddles in interfluvia

To test the hypothesis that land conversion is associated with an increase in the density of ponds and puddles in interfluvia, we as above defined one plateau transect in each of three forests (Seringal, AU and A1 in Tanguro Ranch), three pastures (INCRA Settlement, Cristo Rei Ranch, Lírio Branco Ranch) and three soybean fields (Alvorada, Mutum and Cascavel) (Fig 1). The focal pasture (INCRA Settlement) and soybean field (Alvorada) received two additional plateau transects each for a total of 13 plateau transects. In February 2012 we recorded the number and dimensions of each water body ≥ 5 cm deep that could be sighted from the transects, while estimating the maximum distance to each side of the transect in which a waterbody would be visible if present.

#### Area of shallow, semi-lentic riparian floodplains

To test the hypothesis that land conversion is associated with an expansion in the area of riparian floodplains, we characterized channel morphometry of three first-order streams in forest (APP 2, APP2A, APPM) and three in soybean fields (Tanguro A, B, and C; Fig 1). No streams were characterized in pastures. Because the putative mechanism for floodplain expansion is increased baseflow, more clearly evidenced by dry season water yields (Heyhoe et al. 2011), channel morphometry was characterized in October 2013. Stream channel characterization was conducted in three 50-m sections at 0, 1000 and 2000 m from the headwaters. For streams in converted land, we also characterized one 50-m section in one reservoir contained within the same distance. Each 50-m section was crossed by 6 transects used to measure channel width and water depths. We used the cut off depth of 10 cm for estimating area of shallow water because this is where our indicator species of lentic riparian waters is most frequent (see below). Note that sampling design for quantifying impoundments and ponds and puddles, and area of shallow riparian floodplains, differed due to constraints imposed by the corresponding biodiversity surveys (below): amphibian calling surveys used in the former require considerably less effort and therefore permit greater replication than fish surveys used in the latter.

### Testing for a land use-driven lentification of the freshwater fauna

#### Consequences of impoundment conditions to fish assemblages

We compared the ichthyofaunas of lotic and lentic (i.e. reservoir) sections in the three soybean streams described in ‘Area of shallow riparian floodplains’. Dipnetting was the only methodology appropriate for fish sampling in the narrow, shallow streams rich in coarse woody debris; low water conductivity prevented electrofishing, and gill and seine netting were impossible. We employed a standardized sampling effort of 100 dipnetting person*minutes in each stream section, with a 30-cm diameter dipnet. Because dipnetting is only effective in water ≤ 1m, fish assemblage characterization in impoundments was complemented with three hauls of a seining net (9 m long, 2 m high, 5-mm mesh) and an overnight set of six nylon gillnets (20 m long, 2 m high; and 3, 4, 5, 6, 7, 8 cm distance between diagonally opposite knots).

#### Consequences of shallow riparian floodplains to fish populations

We assessed the relationship between area of shallow riparian floodplains and the abundance of lentic fishes by sampling *Melanorivulus megaroni* (Cyprinodontiformes, Rivulidae) in water < 10 cm deep in all lotic and lentic stream sections described in ‘Area of shallow riparian floodplains’. *M. megaroni* was selected as an indicator of semi-lentic riparian habitat for occurring almost exclusively in shallow margins of streams, swamps and lakes [19]. Preliminary surveys indicated that in marshy floodplains and streamside ponds *M. megaroni* is the dominant fish species, occurring in flooded areas so shallow that one can barely see free-standing water amid the leaf-litter (see also 20 for data on habitat use by rivulids in similar habitats). All fish were euthanized with benzocaine, preserved in formalin 10%, and stored in ethanol 70%.

#### Consequences of land use to amphibian assemblages

We conducted calling surveys at six sampling stations (0, 400, 800, 1200, 1600 and 2000 m) along each of the 26 transects described above. Each transect was sampled between 19:30h and 01:00 (± 30 min) on three dates randomly distributed over January and February 2012. Upon arrival at a sampling station we recorded, for a period of three minutes, calling amphibian species in six categories of abundance (0=no individuals; I=1-2; II=3-5; III=6-10; IV=11-20; V=21-50; VI=50+ individuals). For each sampling station we defined richness as the number of species recorded across all sampling dates, and the abundance of each species as the highest category of abundance recorded. We calculated a conservative index of aggregated (i.e. total) amphibian abundance in each sampling station as the summation of the number of species in a category of abundance multiplied by its lower abundance bound [(N_spp_ in category I * 1) ± (N_spp_ in category II * 3) + …].

## Data analysis

All statistical analyses followed the rationale of testing a given effect by comparing nested linear models [21]. For instance, to test the hypothesis that the expected value of a given response variable differs among land uses (besides all other putative effects), we compared with a log-likelihood ratio test (LRT) two linear models that differed only by the inclusion of land use as a fixed effect. An increase in fit was gauged by the Deviance (D), which is twice the likelihood ratio between the two models, and which converges to a Chi-square distribution under the null hypothesis of no improvement in fit [22]. We used Poisson or Negative Binomial models for count data that had small values, and Gaussian models otherwise. In many cases we used mixed-effects models [23] to take into account the dependence among sampling units (e.g. sampling stations within the same transect). Compliance of fitted models to their assumptions (e.g. constant variance and normality at the link scale, residuals with zero mean and without non-linear trends) where checked from the analysis of the appropriate residuals. All analyses were done in R 3.3.2 [24], with the additional packages lme4 [25] and DHARMa [26]. Expected values and associated standard errors for mixed-effect models were estimated by model-based parametric bootstrap with the function bootMer from the lme4 package [25].

We tested a lentification effect of land use by comparing LRT models that fit different expected values of ponds and puddles/transect, numbers of dams/transect, or areas of shallow riparian floodplain/stream section, with null models that fit the same expected value in all transects or stream sections. We used Poisson glm in the first two cases and a mixed-effect Gaussian glm with stream as a random effect in the third case, to take into account the variance shared by sections within the same stream.

Restricting our analysis only to the dominant fish species in the assemblage (relative abundance >2% [27]), we used LRT to compare mixed-effect Gaussian glm fitted to the log of catch per-unit-effort (CPUE) of each species in each stream section. The first model had only the additive fixed effects of fish species and habitat type (i.e. lotic versus lentic); the alternative model had an additional species-by-habitat interaction term. To test the fixed effects of area of shallow riparian floodplain and land use on *M. megaroni* abundance we used a Negative Binomial glmm because preliminary analyses indicated aggregated distribution of fish counts over sampling units. In both cases stream was a random effect.

We evaluated the effects of lentification on richness and aggregated abundance of anurans recorded at each sampling station. For streamside transects we used LRT to compare mixed-effects models with no fixed effect, the effect of land use, and the additive effects of land use and density of dams in a 800-m radius. For plateau transects we compared mixed-effects models with no fixed effect, the effect of land use, and the additive effects of land use and density of puddles in the transect. We used Poisson models for richness and Gaussian models for the logarithm of the abundance index. All models included transect as a random effect to take into account the variance shared by stations within the same transect.

## Results

### Testing for a land use-driven lentification of the landscape: freshwater habitat change

Number of dams along stream transects increased from zero in forests to a median of 0.4 in pastures and soybean fields (D=11.3, 2 df, p=0.035; Fig 2A). Area of shallow riparian floodplains roughly doubled in soybean watersheds (average ± 1SD = 118 ± 70 m^2^, range 25-246 m^2^) relative to forested watersheds (58 ± 53 m^2^, range 0-142 m^2^; D=4.3, 1 df, p=0.037; Fig 2B). Density of ponds and puddles increased from zero in forest plateaus to a median of 3/ha in pasture plateaus and 2/ha in soybean plateaus (D=54.9, 1 df, p<0.001; Fig 2C; see S1 Figure and S2 Figure for pictures).

**Fig 2.**
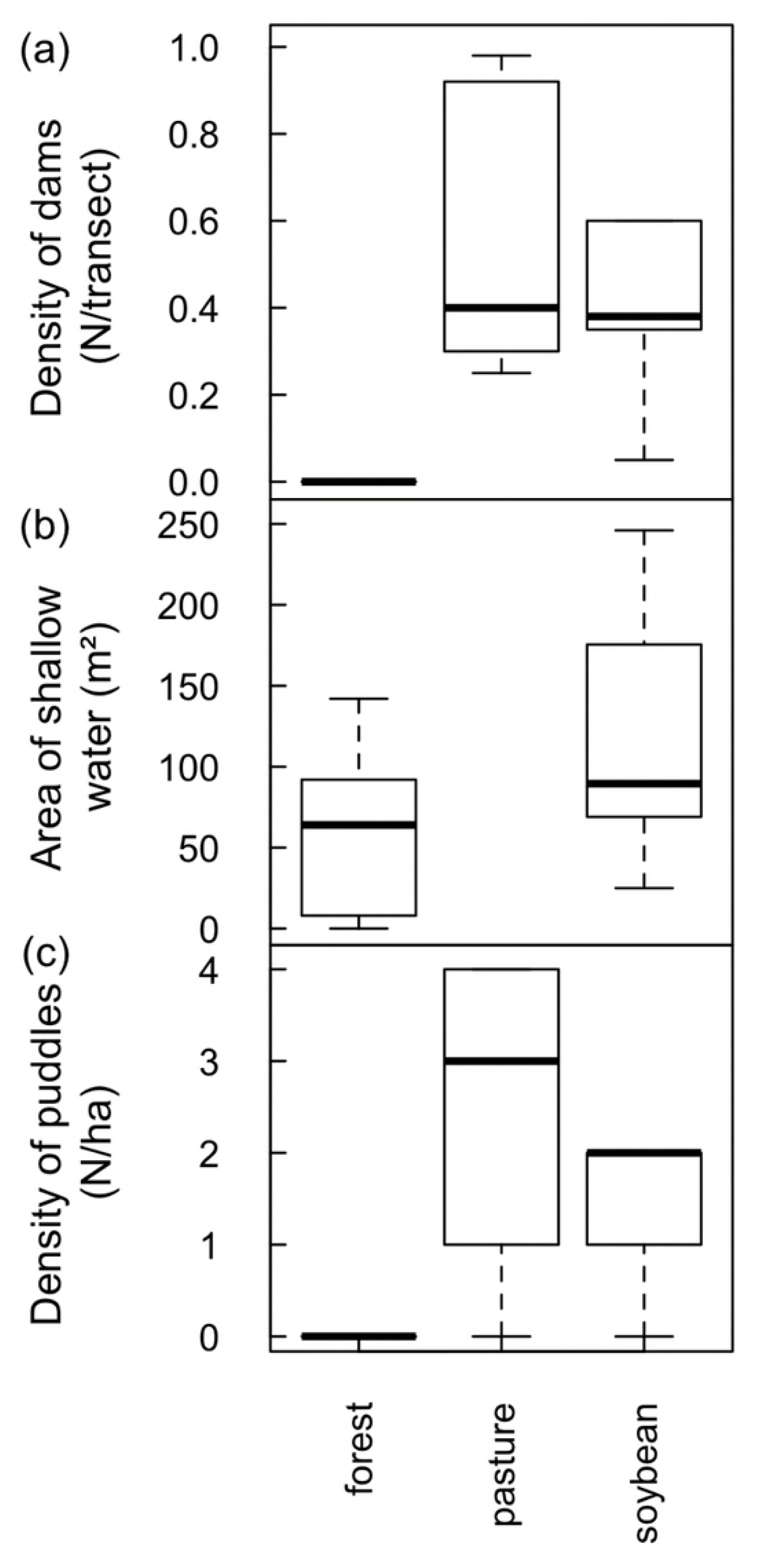
Land use-driven lentification of Southern Amazonian landscapes. (a) Density of dams along streamside transects (b) Area of shallow water in riparian floodplains and (c) Density of ponds and puddles along plateau transects. In (b) note that no streams were sampled in pasture watersheds but that in (a) and (c) no reservoirs and puddles were found along streams and in plateaus, respectively, covered by primary forest.

### Consequences of reservoir formation to fish assemblages

Dominant species in the fish assemblage were *Melanorivulus megaroni* (Rivulidae; average relative abundance across all 12 stream sections 48.3%), *Pyrrhulina australis* (Lebiasinidae; 14.6%), *Aequidens michaeli* (Cichlidae; 11.8%), and the characids *Hyphessobrycon mutabilis* (8.0%), *Astyanax multidens* (7.8%) and *Moenkhausia phaeonota* (2.9%).

These six species, which together comprised 93.8% of all fish individuals dipnetted in the three converted streams, responded strongly to reservoir conditions. CPUE was higher in reservoirs for *Aequidens michaeli, Pyrrhulina australis* and *Astyanax multidens*, but lower for the three remaining species (Fig 3). A significant habitat-by-species interaction term (D=20.5, 5 df, p=0.001) confirmed that some fish species were favored while others disfavored by stream impoundment. In fact, *Hyphessobrycon mutabilis* and *Moenkhausia phaeonota* were entirely absent from reservoirs, as were another ten less common species (< 0.5% in relative abundance). Nine fish species were found exclusively in reservoirs (S1 Table).

**Fig 3.**
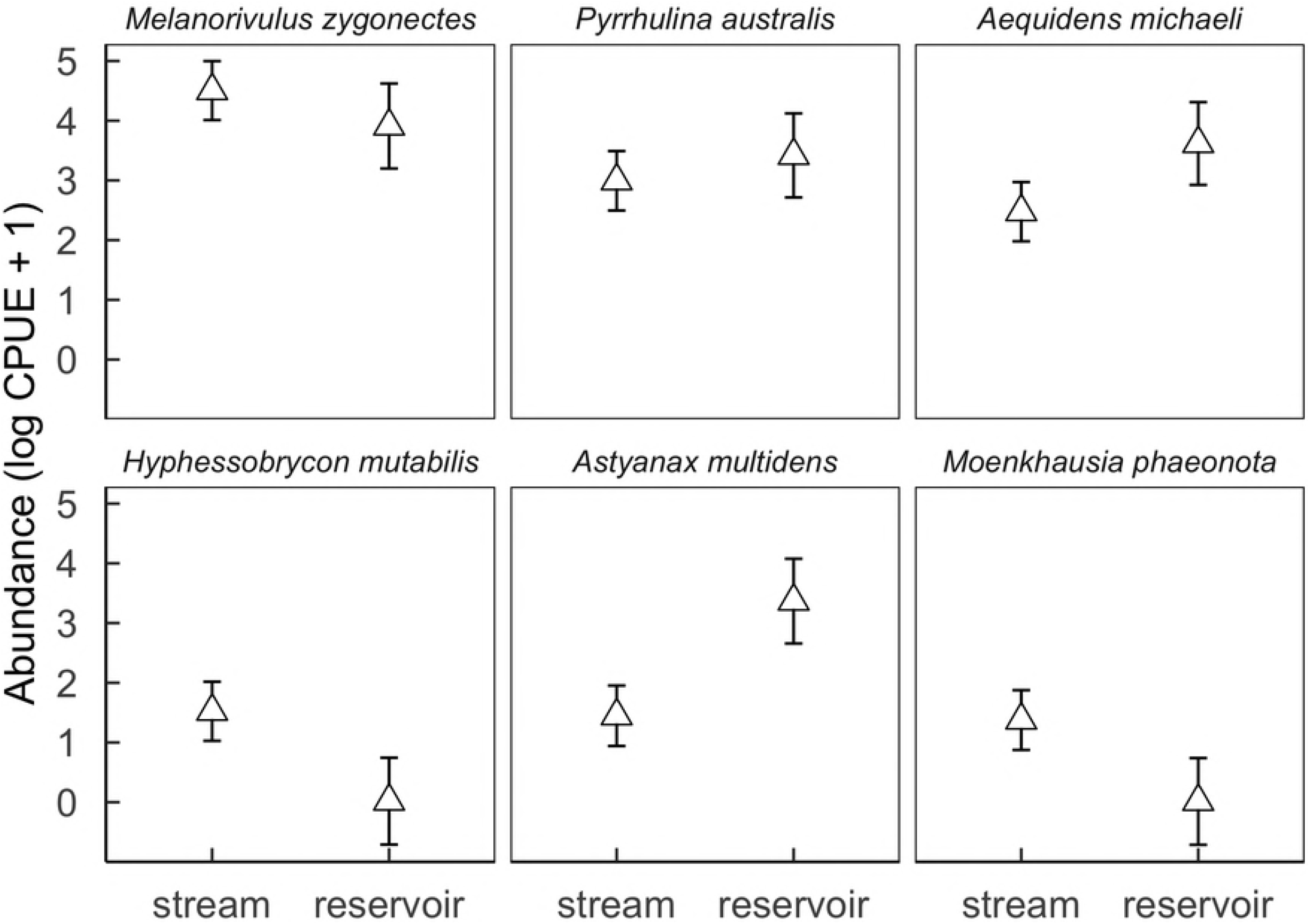
Effect of habitat (stream vs reservoir) on the abundances of the six dominant fish species in streams of deforested (i.e. soybean) watersheds. Symbols and vertical bars represent, respectively, observed abundance means (ln CPUE + 1) ± 1 standard errors of the estimated values by a linear mixed effect Gaussian model, accounting for all sources of variation (fixed and random effects).

### Consequences of floodplain expansion to the abundance of *Melanorivulus megaroni*

*M. megaroni* abundance was positively related to the area of shallow riparian floodplains (D=17.2, 1 df, p<0.001; Fig 4).

**Fig 4.**
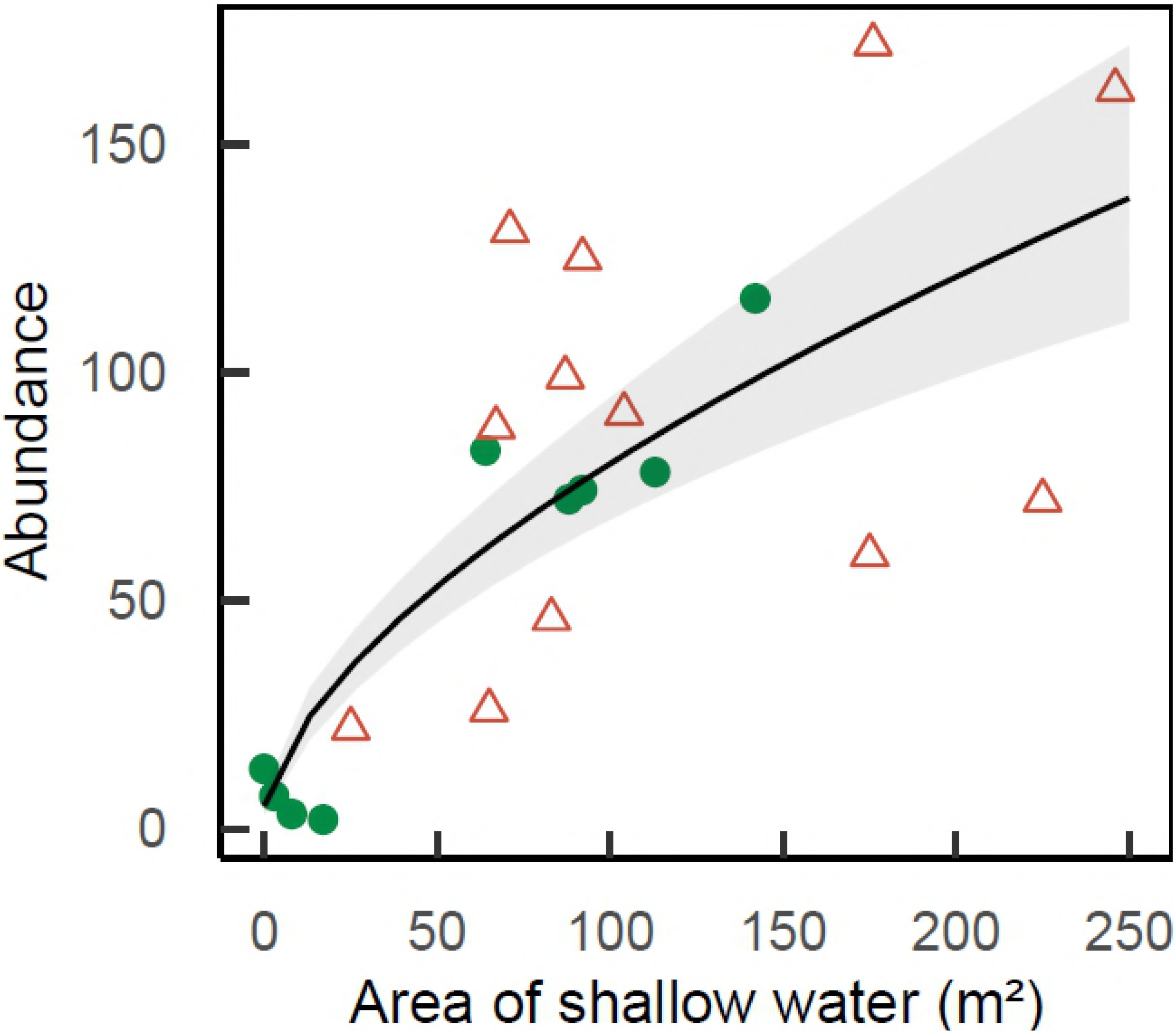
Abundance of *Melanorivulus megaroni* as a function of the area of shallow water in each 50-m stream section (green solid circles: streams in forested watersheds, red open triangles: streams and reservoirs in deforested watersheds). The line is that predicted by a negative binomial generalized model with random effects due to streams. The grey area shows ± 1 the standard errors of the estimated values by the model, accounting for all sources of variation (fixed and random effects).

### Consequences of land use to amphibian assemblages along streams

Transects in deforested watersheds had more than twice as many calling species (20 species) than transects in forested watersheds (9 species). Note, however, that rarefaction curves asymptoted in pastures and to a lesser degree in soybean fields, but not in forests (Fig 5A). Because more singletons and doubletons were found in forest transects, Chao2 estimators of species richness were 28.8 ± 16.3 for forests (mean ± 1 SD) but 20.3 ± 0.9 for pastures and 21.6 ± 2.1 for soybean fields.

**Fig 5.**
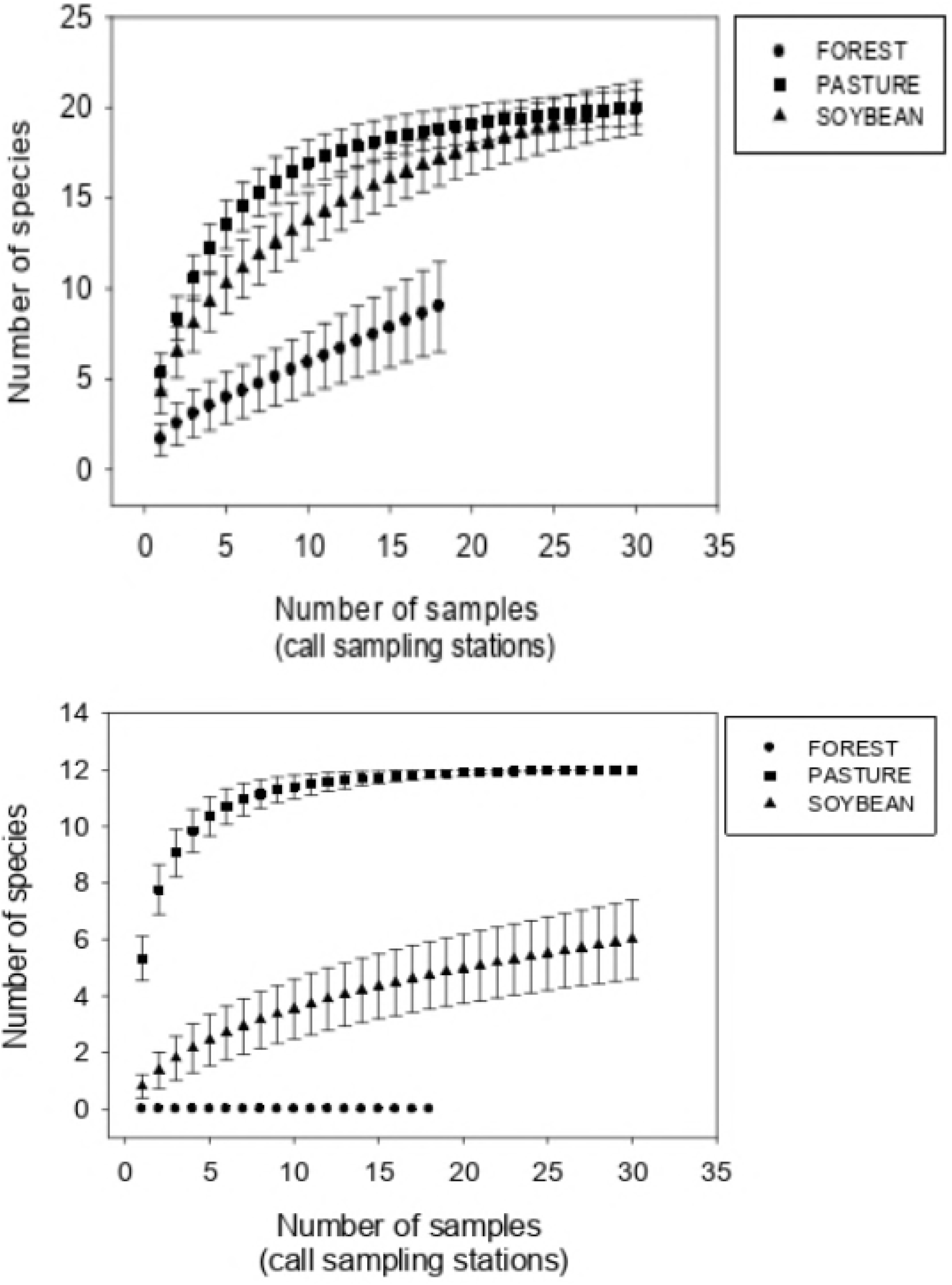
Rarefaction curves of amphibian richness along (A) streamside transects and (B) plateau transects in forests, pastures and soybean fields. Symbols represent estimated means ± 1 SD. Note that there were three streamside and three plateau transects in forests (therefore 18 call sampling stations) but five streamside and five plateau transects in pastures and soybean fields (therefore 30 call sampling stations). Note also that in B no amphibian calls were recorded in forests, where ponds and puddles are absent and where water bodies are invariably associated with streams and their floodplains.

No species was heard exclusively along streams in forest watersheds although the only sightings of *Phyllomedusa vaillanti* (Hylidae) and *Leptodactylus pentadactylus* (Leptodactylidae) occurred in those streams. By contrast, 20 species were heard exclusively along streams in deforested watersheds (S2 Table).

Amphibian species richness per sampling station increased in converted land relative to forests (D=12.4, 2 df, p=0.002), and with number of dams within an 800-m radius (D=7.3, 1 df, p=0.007). Predicted values confirm a much higher species richness in pastures and soybean fields than in forests (Fig 6A). Amphibian aggregate abundances per sampling station also increased in converted land relative to forests (D=14.2, 2 df, p=0.001), but the effect of number of dams was weaker (D=3.5, 1 df, p=0.06). Still, we used the model with both predictors to estimate the expected values, which again confirmed higher abundances in pastures and soybean fields relative to forests (Fig 6B).

**Fig 6.**
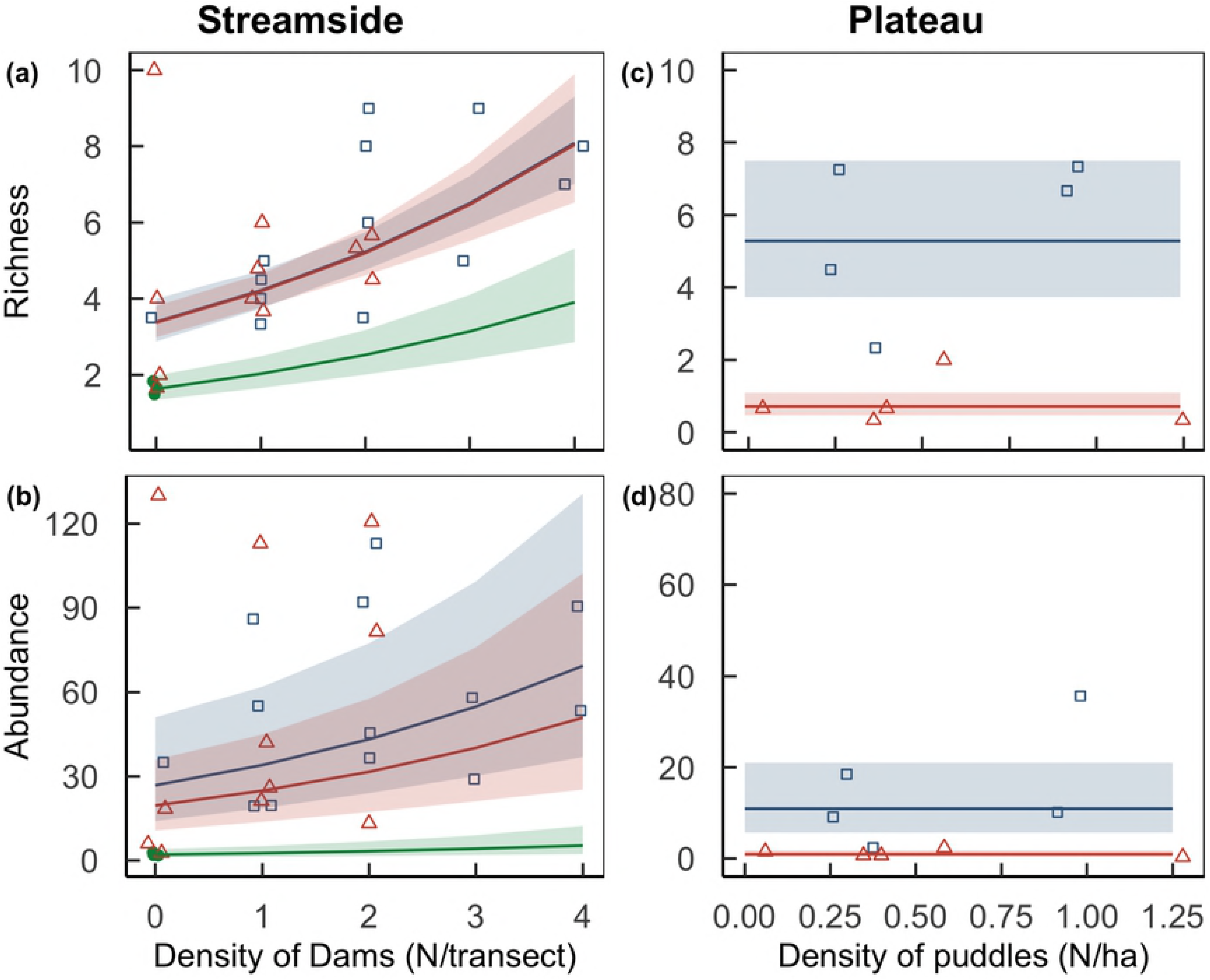
Amphibian richness (upper row) and aggregate abundance (lower row) per sampling station as a function of the density of dams along streams (left column) and puddles in plateaus (right column) in forests (solid green circles), pastures (open blue squares) and soybean fields (open red triangles). Curves represent trends (± 1 standard errors of estimated values) predicted by the best supported model (generalized linear models with random effects of the transects). Note that in plateaus only land use had a significant effect on amphibian abundance and richness, which is depicted by the horizontal lines of predicted values and standard errors. In primary forest plateau transects no amphibian species were recorded, and the model was fitted only to data gathered in soybean and pastures. Points are jittered along x and y axes to improve data visualization.

### Consequences of land use to amphibian assemblages in plateaus

Land use had an even stronger effect on amphibian assemblages in plateaus. Transects in forest plateaus were entirely devoid of calling amphibians. By strong contrast, pastures and soybean fields had respectively 12 and 6 calling species (S2 Table, Fig 5B). Rarefaction curves asymptoted in pastures but not in soybean fields; Chao2 estimators of species richness therefore coincided with measured richness in pastures (12.0 ± 0 species) but not in soybean fields (8.9 ± 4.3 species).

Because forest plateaus were devoid of calling amphibians, we could not fit models to predict amphibian richness or abundances for all three land use types considered jointly. Considering converted plateaus only, both amphibian richness (D=15.3, 1 df, p<0.001; Fig 6C) and aggregated abundance per sampling station (D=12.2, 1 df, p<0.001; Fig 6D) was higher in pastures than in soybean fields. In both cases predicted values were higher in pastures than in soybean fields. No improvement of fit was obtained by including pond and puddle density.

## Discussion

This study provides evidence, for the first time, that deforestation promotes an increase in the availability of lentic freshwater habitats in Southern Amazonian landscapes. Streams in converted watersheds had numerous impoundments, and riparian floodplains twice as wide than streams in preserved watersheds. At the same time, converted interfluvia had numerous ponds and puddles, in a striking contrast to the complete absence of waterbodies in forested interfluvia. Importantly, lentification of streams and interfluvia was accompanied by strong responses by fish and amphibian assemblages.

Reservoir conditions favored half of the six dominant fish species, and disfavored the other half. The characid *Astyanax multidens* and the cichlid *Aequidens michaeli* were 8 and 2.3-fold more abundant in reservoirs than in lotic reaches; several other less common cichlids were also predominantly or exclusively found in reservoirs, as were the trahira *Hoplias malabaricus* and the curimatid *Steindachnerina* sp. At the other extreme, not a single individual of characids *Hyphessobrycon mutabilis* and *Moenkhausia phaeonota*, otherwise common in stream reaches, was found in reservoirs. These patterns are consistent with large-scale studies showing that cichlids, characids and other characiforms, but also erithrynids, serrasalmids, loricariids and curimatids that inhabit more lentic habitats such as backwaters, shallow river margins and lakes, are sometimes favored by river damming [28].

Expanded stream floodplains, in turn, had a strong positive effect on the abundance of *Melanorivulus megaroni*, a fish species associated with shallow waters (>90% of individuals were sampled in water <10 cm deep) and streamside pools. Other fishes could be influenced by floodplain expansion as well given that many small-sized species, or larvae and juveniles of larger species, rely on these habitats as nurseries and shelters against predation and water dragging [29].

Conversion of forests to pastures and soybean fields had a surprising positive effect on streamside amphibian abundance and richness. Both response variables were associated with number of dams around calling sampling stations. Thus, landscape lentification via reservoir formation is a plausible contributor to this phenomenon, although other hydrological and non-hydrological changes associated with deforestation, including the expansion of stream floodplains (not quantified along surveyed streamside transects) and canopy opening, could also be involved.

Even more striking was the influence of land conversion in interfluvia on amphibian richness and abundance. No single amphibian species was heard in forested plateaus, as opposed to 12 in pastures and six in soybean fields. Because calling involves not only benefits (mate attraction and selection) but also costs (energetic costs, increased risk of predation and parasitism), males usually call in, at or close to their breeding sites [30]. Therefore, the lack of water bodies is a sufficient explanation for the absence of calling amphibians in forested plateaus, as 35 of 38 amphibian species in the region depend on water for egg laying and/or premetamorphic development [31]. We suspect the lack of a statistical effect of density of ponds and puddles on amphibian richness and abundance results primarily from the need to exclude forested plateaus from the analysis, and secondarily from a mismatch in the scale of habitat assessment (ponds that could be seen from transect, i.e., tens of meters) and that of faunal assessment (frogs that could be heard from transect, i.e., hundreds of meters).

Contrary to streams, where amphibian richness and abundance were similar in pastures and soybean fields, in interfluvia pastures had twice as many amphibians as soybean fields. At least two factors could be involved. First, several of the waterbodies in pastures were cattle ponds, which, contrary to roadside puddles, are constructed to effectively hold water – i.e., they are deeper and compacted with tractors. As a consequence, waterbodies in the focal pasture held water for one month longer than in the focal soybean field (personal observation), a difference in hydroperiod of considerable biological significance considering that a primary cause of mortality in temporary ponds is dessication [15]. Second, environmental conditions in plateaus - effectively contrasting primary forests, pastures and soybean fields - are much harsher than along streams, at the time of sampling protected by law by a 25-m wide vegetated buffer zone. Specifically, the intensive management of soybean plantations inflicts strong, periodical disturbance to the fauna that inhabits, visits or traverses interfluvia. Conversion of pastures to soybean fields is made by clearing with fire, tilling and liming, whereas plantation management annually involves sowing, applying fertilizers (~300 kg NPK/ha) and pesticides (13.7 kg/ha including 39 active ingredients) and harvesting [3; 32]. Considering that puddles in soybean fields are subject to direct pesticide overspraying and runoff, it is surprising that there are amphibian species colonizing ponds in plantations – although, we do not know whether these are viable habitats for metamorph production or simply amphibian population sinks (as is the case for aquatic insects; see 33).

In summary, the creation of new reproductive habitat is a plausible hypothesis for the increased amphibian abundance in converted land. The increase in species richness is, by contrast, unexpected. Differences in observed gamma diversity are likely artifacts of abundance; Chao estimators predict 29 species in forests, less than the 38 recorded [31] but more than the 22 predicted in converted land. Differences in alpha diversity, in turn, do not appear to be artifacts and could be due to the particular biogeographical scenario of our study site and/or the spread of generalistic open area species with deforestation (see also 34). Our study site is at the edge of Amazonian closed-canopy forests; the savanna line is as close as ~20km south and it is likely that in the Pleistocene this area has alternated open-canopy/closed-canopy phytophysiognomies [35]. The observation that several species found in closed-canopy forests are more typical of open areas (*Leptodactylus labirinthicus, L. mystaceus, Hypsiboas albopunctatus, Rhinella schneideri*) is a testimony for this mixed amphibian assemblage. On top of that, converted streamside transects have the addition of at least seven species exclusive of open areas *(Leptodactylus fuscus, Eupemphix nattereri, Physalaemus cuvieri, P. centralis, Pseudopaludicola mystacalis, Scinax fuscovarius* and S. *fuscomarginatus).* Converted watersheds could therefore be housing an enriched amphibian fauna through the mixture of closed-canopy species of Amazonian origin (e.g., *Osteocephalus taurinus, Phyllomedusa vaillanti, Pristimantis fenestratus*), open-area cerrado species capable of colonizing closed-canopy forests, and generalistic open-area cerrado species that accompany the line of deforestation and are favored by landscape lentification. Of course, at this point it is impossible to know whether the increased species richness in converted land is a stable endpoint or a product of transient dynamics given the recent deforestation in the area (< 40 years), the even more recent decrease of matrix quality via conversion of pastures to soybean fields (< 10 years), and the presence of relatively large fragments of primary forest in the region and riparian buffer zones (Fig 1).

### Broader geographic consequences

Although our research is local, it is tempting to speculate that the lentification of freshwater systems and their biota could be occurring in many locations of deforested Amazonia. This is because all proposed drivers of lentification are quite general. We reviewed the available evidence linking land conversion and the density of dams, soil compaction and stream discharges in the Amazon Basin (and its sub-basins Xingu, Tapajós, Madeira, Solimões, Negro and Amazonas) and surrounding basins (Tocantins, Marajó and the NW Coastal South Atlantic to the Southeast; Orinoco and Coastal North Atlantic to the North) (Table 1). A total of 179 hydroelectric dams are currently operating or under construction, and another 275 planned, in four basins (Tocantins, Amazon, Orinoco, Coastal North Atlantic), including all six Amazon sub-basins [12; 35]. Dramatically increased densities of small impoundments in deforested land were reported in the Xingu^6^ and the Amazonas sub-basins [36].

**Table 1.**
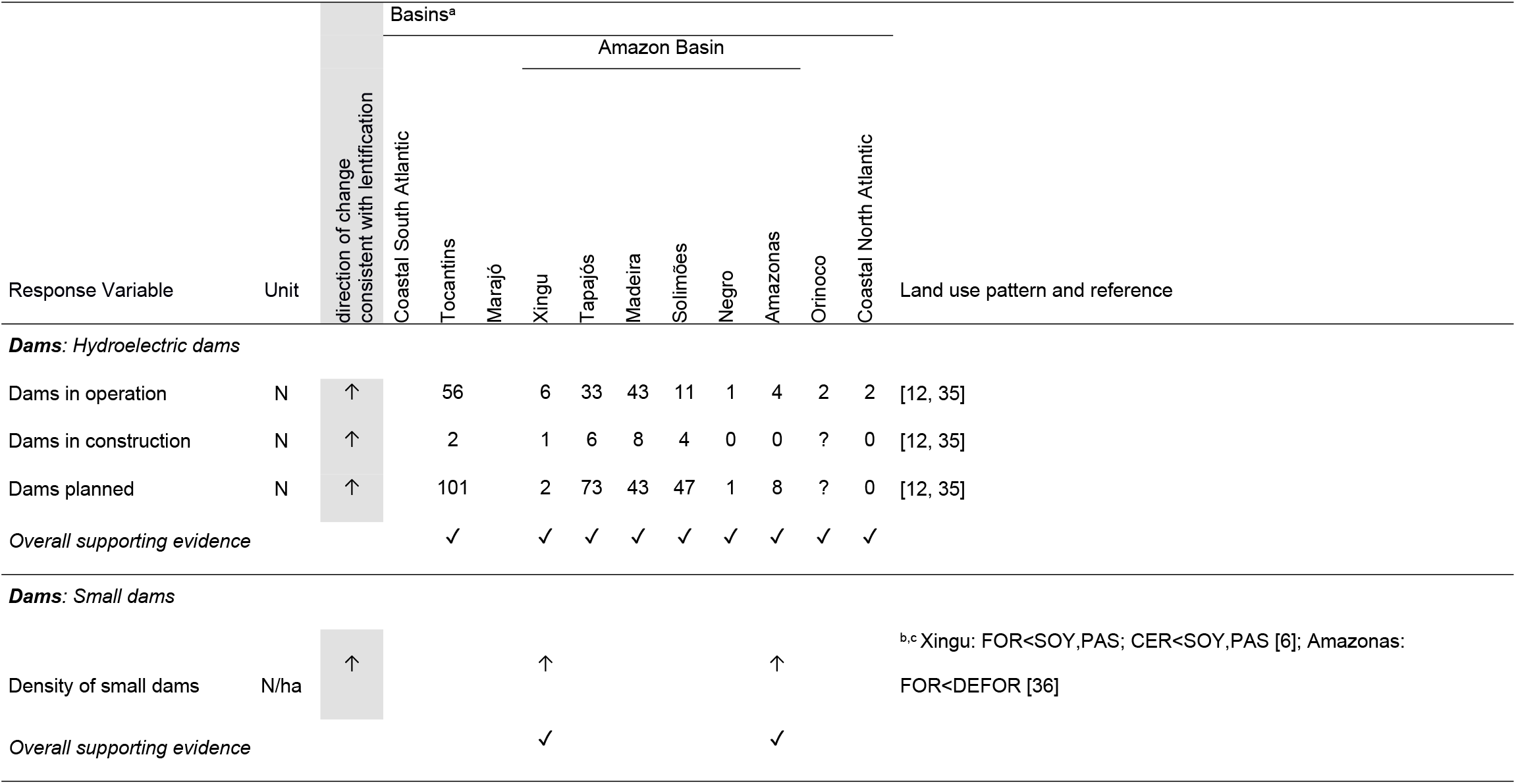
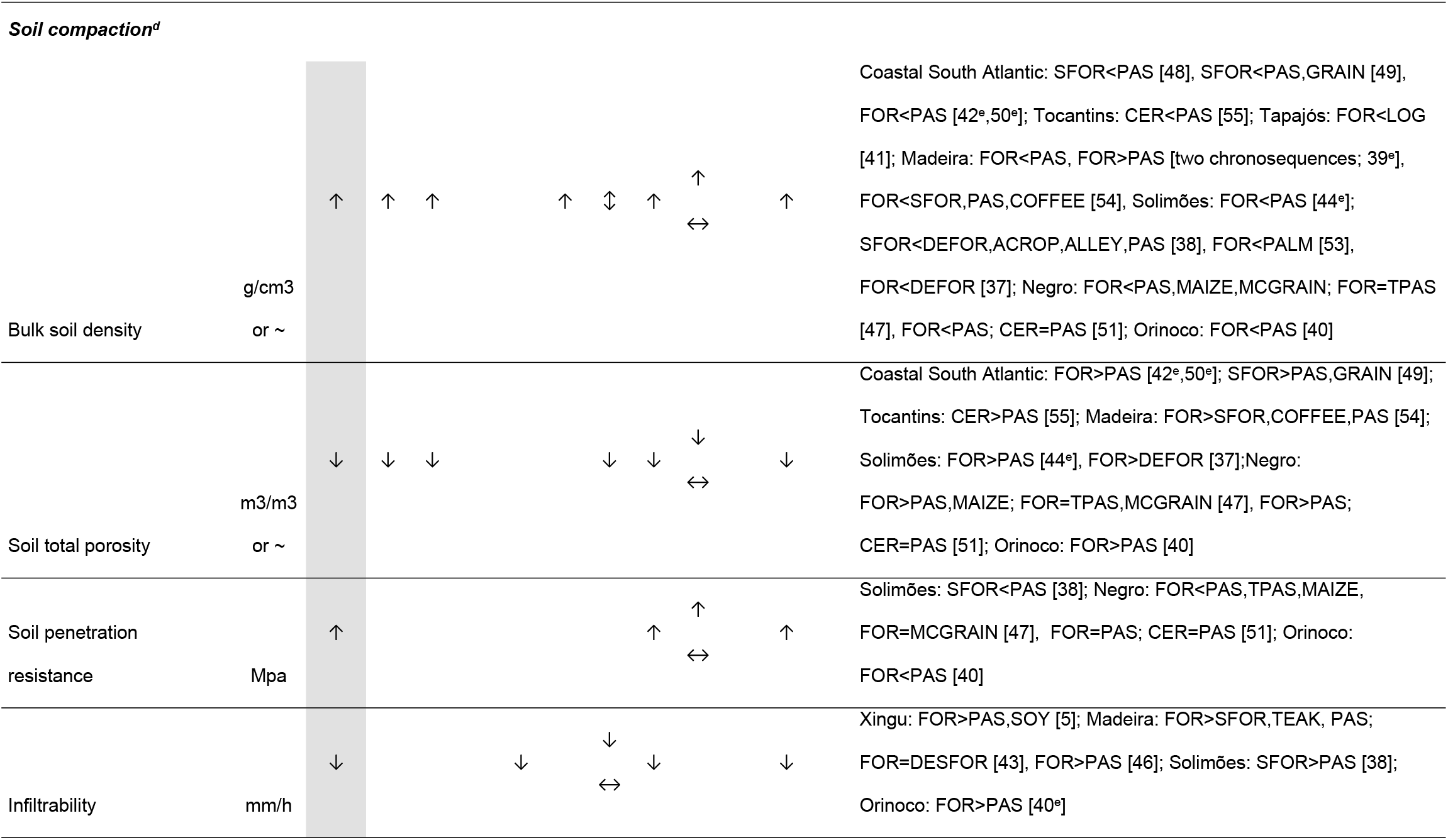
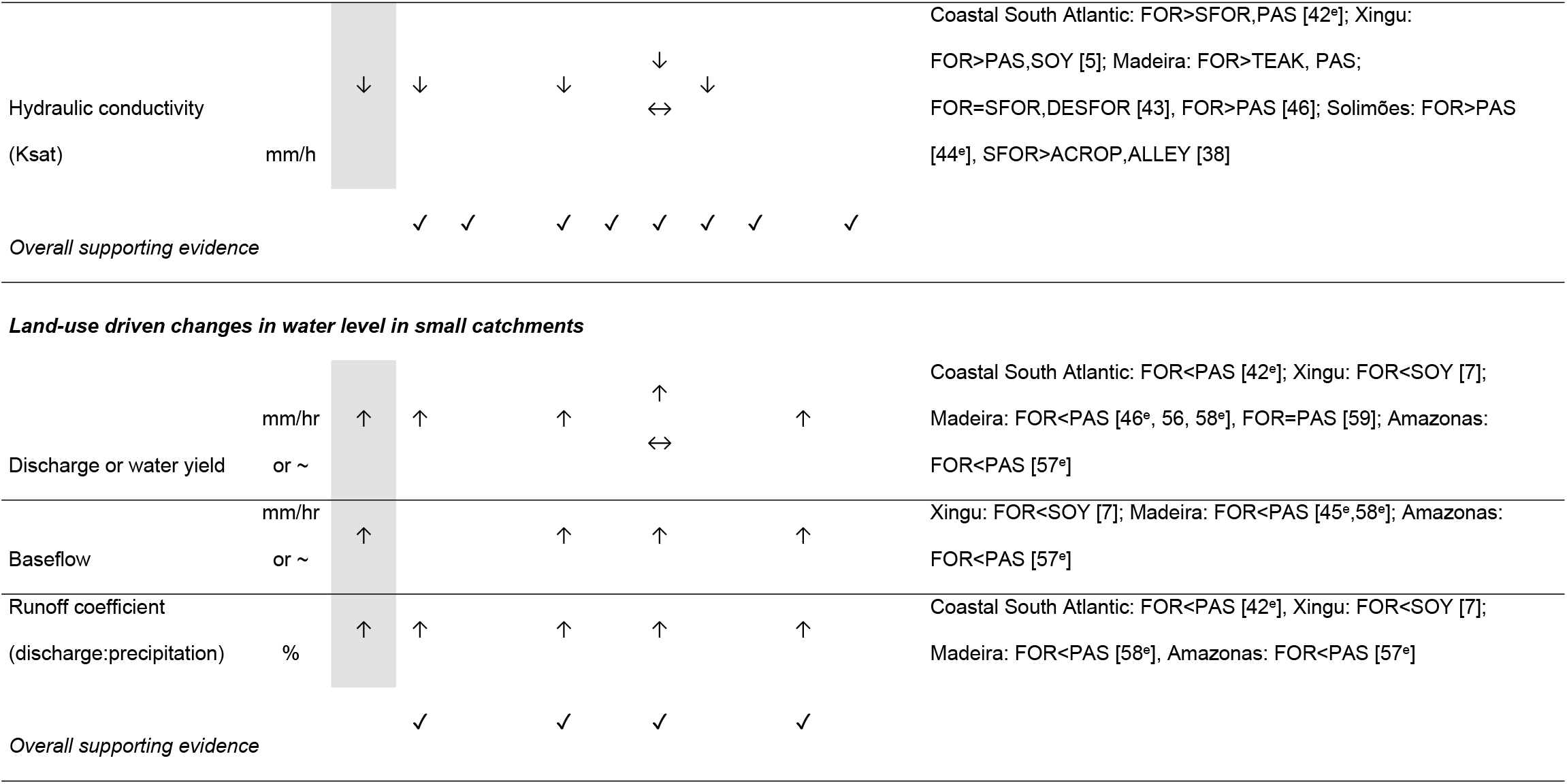
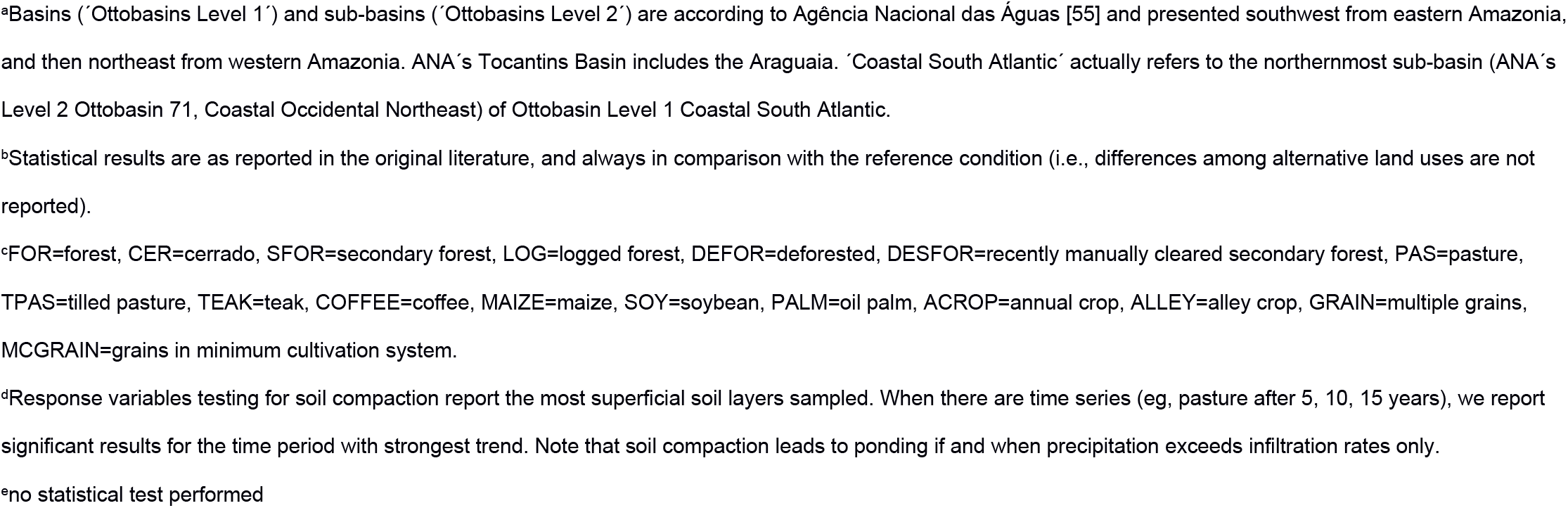
Land-use driven changes in landscape hydrological properties - how widespread are drivers of lentification in deforested Amazonia? Arrows represent an increasing trend (↑), a decreasing trend (↓), both increasing and decreasing trends (↕) and no trend (↔).

Soil compaction, as indicated by increased bulk soil density or soil penetration resistance, and/or by decreased soil porosity, infiltrability or hydraulic conductivity, were consistently observed in all 19 studies we could find conducted in four basins (Coastal South Atlantic, Tocantins, Amazon, Orinoco), including five of six Amazon sub-basins (Xingu, Tapajós, Madeira, Solimões and Negro) [37; 38; 39; 40; 41; 42; 43; 44; 45; 46; 47; 48; 5; 49; 50; 51; 52; 53; 54; 55] (Table 1). Importantly, soil compaction was observed in every type of land use including secondary and logged forests, deforested land, pastures, and in soybean, maize, oil palm, coffee, and teak plantations (Table 1). Among them, it is precisely in pastures – by far the prevailing land use in converted Amazonia - where soil compaction is most intense. Relative to forests, infiltrability in pastures decreased from 8 to 162 times depending on factors such as soil type, pasture age, and cattle stocking densities. Even though soil compaction only results in ponding if precipitation exceeds infiltration rates, increased overland flows in pastures are commonly reported. In eastern Amazonia rainfall rarely if ever exceeds infiltration capacities in forests; in pastures, by contrast, around ¾ of storm-averaged rainfall intensities exceed infiltration capacities [42].

Finally, because deforestation strongly reduces evapotranspiration, a larger fraction of rainfall is exported from the watershed as streamflow. This pattern, commonly observed in small catchment studies in both temperate and tropical countries [56], was recorded in two basins (Coastal South Atlantic and Amazon), and three of the Amazon sub-basins (Xingu, Madeira and Amazonas) [42; 56; 57; 58; 45; 7]. Conversion of forests to pastures and soybean fields increased water yields, discharge and baseflow, and discharge-to-precipitation ratios (Table 1). Interestingly, one study that failed to observe a linkage between land conversion and discharge nevertheless observed stream structural changes consistent with lentification. Deegan et al. [59] found that in the conversion of forests to pastures the pool-and-run structure was replaced by a narrow run of open water bordered by wide marshy areas invaded by riparian grasses (mean wetted width was 5X greater in pastures than forests, N=18 streams). Ground-truthed remote sensing permitted the authors to infer that the infilling of stream channels by riparian grasses, which slows down water velocities and promotes the transient storage of water, occurs in almost all of the small pasture streams in the Ji-Paraná Basin, a 74,000 square kilometer sub-watershed of the Madeira [59].

### Concluding remarks

In this study we demonstrate that extensive hydrological change occurs very early in the process of frontier settlement. The deliberate construction of reservoirs, cattle and drainage ponds adds to the unintentional multiplication of puddles and expansion of floodplains to modify existing habitat, or even create new habitat, for freshwater organisms. The phenomenon of freshwater lentification we here propose, if indeed shown to be widespread in deforested Amazonia, could have significant consequences to biodiversity and human health; the dramatic upsurge in malaria and schistosomiasis following deforestation, for example, is partly a consequence of increased availability of lentic freshwater habitats in the landscape [60,61].

It is reasonable to expect that other drivers of lentification not considered in this study could be currently operating or become important in the future. For example, reservoir management may promote the rise of the water table [62], potentially increasing the area of river floodplains. By contrast, unsustainable water extraction, large scale deforestation [14] or climate change could all contribute to lowering water tables, reducing floodplain widths and causing flow intermittency and increased stream temporality. Climate change and anomalous climatic events, in particular, could strongly interact with other phenomena described in this article to magnify the intensity of freshwater lentification. Such interactions could be particularly strong in the Amazonian Arc of Deforestation, which is not only a region of widespread land conversion – and therefore widespread river damming, change in stream discharges, soil compaction and creation of farm ponds - but also predicted to be highly vulnerable to savannization [14].

## Acknowledgments

We thank Grupo Amaggi (Fazenda Tanguro) and neighboring farmers (Assentamento do INCRA, Fazenda Cristo Rei, Fazenda Lírio Branco) for granting access to their lands, and the Instituto de Pesquisa Ambiental da Amazônia – IPAM (especially O. Carvalho Junior, W. Rocha, P. Brando and M. Macedo), Grupo Amaggi (especially W. De Ré) and M. Rosa (Secretaria da Agricultura e Meio Ambiente de Canarana) for logistical support. G. Cordeiro Junior, G. Cordeiro, V. Dimitrov, I. Calado, D. Nunes, S. Rocha e R. Quintino provided invaluable help in fieldwork, and I. Calado and A. C. Machado in labwork. P. Lefebvre kindly produced the map and M. Leite helped in the statistical analysis. ICMBio provided collection permits (ICMBio 17559). We thank FAPESP (2008/57939-9, 2010/52321-7 to L.S.), CNPq (563075/2010-4 to L.S.) and the International Finance Corporation - Biodiversity and Agricultural Commodities Program for funding this research (to L.S.and B.G.), FAPESP (2011/20458-6 to P.I., 2010/19427-6 to D.D.B.N.) and CNPq for fellowships (Research Grant to P.I.P.), and the University of Veterinary Medicine of Vienna for a travel grant (B.G.).

## Supporting Information Captions

**S1 Figure.** Streams in forested (a) and deforested (b,c) watersheds. (c) is a man-made reservoir. Pictures by Paulo Ilha.

**S2 Figure.** Ponds and puddles in plateaus converted to pastures (a-d) and soybean plantations (e-g). No ponds or puddles were found in forested plateaus. (a) and (b) are man-made cattle ponds; all other puddles (c-g) appear to be formed solely by soil compaction by cattle trampling, road construction and machinery traffic. In the rainy season ponds and puddles like these were inhabited by up to 5 and 3 species of amphibians in pastures and soybean fields, respectively. Those in pastures contained in addition high abundances of predatory insects including dragonflies, bugs and beetles. Pictures by Luis Schiesari and Victor Dimitrov.

**S1 Table.** Pooled number of fishes sampled in lotic and lentic (i.e. reservoir) stream sections in three streams in deforested watersheds, ranked by abundance.

**S2 Table.** Incidence matrix of amphibian species recorded in streamside and plateau transects in forests, pastures and soybean fields (1= calling, ±=observed).

